# Evaluation of network inference algorithms for derivation of metabolic relationships from lipidomic and metabolomic datasets

**DOI:** 10.1101/2025.05.23.654112

**Authors:** Finn Archinuk, Haley Greenyer, Ulrike Stege, Steffany A.L Bennett, Miroslava Cuperlovic-Culf, Hosna Jabbari

## Abstract

**Motivation:** Various methods have been proposed to construct metabolic networks from metabolomic data; however, small sample sizes, multiple confounding factors, the presence of indirect interactions as well as randomness in metabolic processes are of major concern.

**Results:** In this study, we benchmark existing algorithms for creating correlation- and regression-based networks of changes in metabolite abundance and we evaluate their performance across different sample sizes of a generative model. Using standard interaction-level tests and network-scale analyses based on centrality scores, we assess how well these methods capture simulated metabolomic networks. Our findings reveal significant challenges in network inference and result interpretation, even when sample sizes are significant and data are the result of computer modeling of metabolic pathways. Despite these limitations, we demonstrate that correlation-based network inference can, to some extent, discriminate between two different metabolic states. This suggests potential utility in distinguishing overarching changes in metabolic processes but not direct pathways in different conditions.

**Availability:** All relevant data is provided at https://github.com/HosnaJabbari/metabolicRelationships

**Supplementary information:** Supplementary data are available at *Bioinformatics Advances*

## Introduction

The metabolome, encompassing all small molecules within a system, can be profiled using various mass spectrometry and nuclear magnetic resonance “omics” techniques that measure molecular concentrations across systems in high throughput [Wishart, 2007]. Advances in these methods have enabled new approaches to understanding disease, increasingly integrating the analysis of individual metabolite abundances into metabolic networks [Wilkins and Trushina, 2018]. Previous studies have formulated metabolomic networks as graphs, where the nodes represent metabolites and the edges connecting them indicating some level of covariation [Liu et al., 2020]. Metabolites may be associated in these graphs through direct or indirect enzymatic reactions, mass action, signaling, or co-expression. These metabolic association patterns are purported to change upon perturbation of the system. Elucidating the alterations in these patterns is increasingly used to infer underlying biological process. The changes in the metabolic networks direct experimental exploration targeting different steps in a molecular pathway identified in network analysis or interrogated to determine changes that result from pathological conditions [Szymanski et al., 2009].

These bioinformatic approaches face the challenges posed by dynamic modifications in interactions due to a change of state, further complicated by the fact that many interactions within constructed networks remain unknown. To address this, a process called “network inference” is commonly used to deduce edges from metabolic concentration data. However, this process is often constrained by the limited number of reference samples available relative to the larger number of metabolites and their potential interactions [Nyamundanda et al., 2013] as well as difficulty in distinguishing between direct and indirect interactions across large concentration ranges with fast change in concentration level. Network inference algorithms (NIAs) often rely on simplifying assumptions, such as linear relationships, steady-state conditions, and sparsity of interactions, to make computations feasible. However, these assumptions often fail to capture the complexities of biological systems, which are dynamic, nonlinear, and subject to experimental noise. Additionally, inferred connections may reflect correlations rather than true causal relationships, emphasizing the need for careful interpretation and experimental validation.

Jahagirdar et al. [2019] provided a foundation for testing NIAs on a simulated dataset. Using a simulated dataset to evaluate NIAs offers several advantages over biological datasets. Simulated datasets provide a ground truth network, allowing for precise and objective assessments of algorithm performance, including identification of errors and quantification of accuracy. In contrast, biological datasets often lack a known underlying interaction network, with indirect often as strong as direct relationships to, reaction based edges, making it difficult to distinguish between true, direct biological interactions and algorithmic artifacts. Moreover, while simulated datasets enable controlled experimentation by systematically varying factors such as sample size, noise levels, and association strengths, this control is not only challenging to achieve with biological data, it may mask real biological processes.

This paper seeks to address these issues by extending the work of Jahagirdar et al. [2019] to better understand how (1) evaluation metrics change with small sample sizes, (2) investigate where errors occur in inferred networks, and (3) establish how inferred networks can be used to differentiate metabolite states. Informed by graph theory, we apply centrality measures that demonstrate how they can be used to identify bottlenecks or other important regions within a network based on the overall network structure [Liu et al., 2020]. We find that network inference based exclusively on pairwise interactions between metabolites fails to accurately capture the biological network with centrality measures clearly demonstrating these limitations. These deficits are not simply due to a lack of samples rather they demonstrate a failure to converge to the reality of the underlying network. This work adds to the growing movement to examine the assumptions baked into pairwise NIAs [Lingjærde et al., 2020, Peel et al., 2022, Li et al., 2024]

## Materials and Methods

### Simulated Metabolic Network

To evaluate NIAs, we first used synthetic data from a simulated network to ensure a known target network. Previous studies have relied on known pathways, but novel connections cannot be definitively classified as an inference error or an undiscovered interaction [Marbach et al., 2012]. Specifically, we used the model developed by Jahagirdar et al. of Arachidonic Acid (AA) degradation and benchmarked this model against several NIAs [Jahagirdar et al., 2019]. The simulated network was constructed based on known reactions involved in AA metabolism. The constructed AA metabolic model included 83 metabolites and 131 reactions. Reactions were written as ordinary differential equations, with enzymatic reactions described using Michaelis-Menten kinetics and non-enzymatic mass action laws [Michaelis et al., 2011]. Figure 1 shows the interactions between metabolites. Edges indicate a reaction in the AA model; removal of metabolites from the system are not shown but are present at each terminal node. The network aggregates multiple metabolites under the umbrella of phosphatidylcholine (PC). This simplification is secondary to the focus of our work of evaluating network inference algorithms against a biologically inspired metabolomic ground truth.

**Fig. 1.**
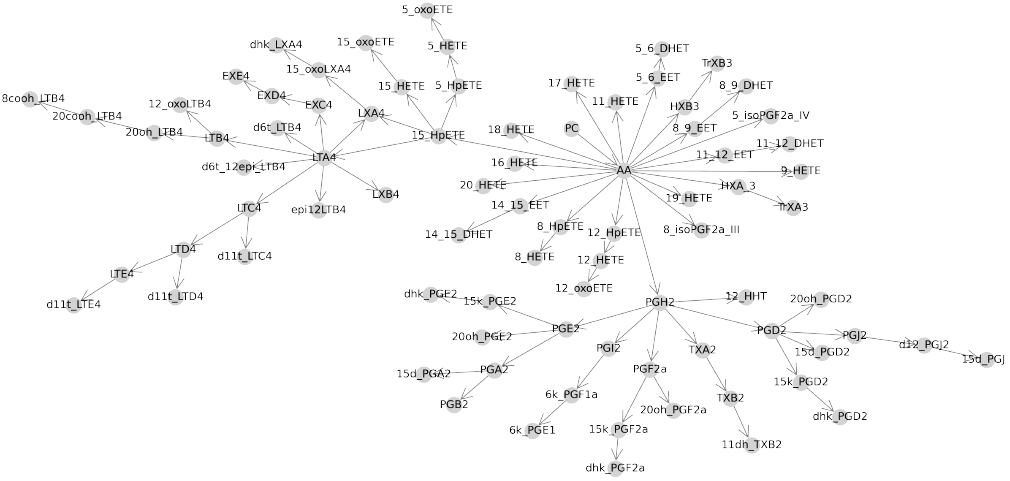
The simulated Arachidonic Acid network representing primary interactions. Nodes represent metabolites. Connections between nodes indicate a direct reactionary interaction, either through enzymatic conversion or mass action.

The adjacency matrix for the AA model provides information about the presence or absence of reactions. This is a *n × n* matrix, where *n* represents the number of nodes (i.e., metabolites) in the network, acting as our ground truth. Interactions between two nodes are indicated by a ‘1’ in the corresponding row and column of the matrix if there exists a reaction, otherwise ‘0’ indicates no interaction. The simulated network generates samples by initializing reaction parameters and running the model forward until steady state is reached. By randomizing initial reaction parameters, we created any number of samples, each in the form of a vector: (*a, b*, …, *z*) where *a, b*, … and *z* are metabolite concentrations measured at the end of the simulation.

### Evaluation Metrics

In its simplest form, all possible pairs of metabolite interactions can be organized into a matrix (*A*) representing all potential interactions. It is assumed that weaker interactions indicate pairs that are more distant in the network. These interactions were quantified using an association function:

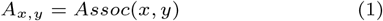

where for each entry *A*_*x*,*y*_ of *A, x* and *y* represents a given pair of metabolites, and *Assoc* represents an association function produced by a given NIA. When a threshold *τ* is applied to the entries of *A*, a binary matrix *B* is obtained:

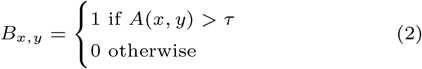

with *τ* being a fixed value, typically 0.6 for correlation networks [Suarez-Diez and Saccenti, 2015]. When applicable, the absolute value is used, particularly in correlation-based methods where strong negative correlations are also considered significant. To remove the influence of tuning a threshold value, we dynamically selected a threshold for each inferred network to maximize the F1-score. The F1-score is calculated as the harmonic mean of precision and recall:

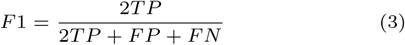

where *TP*, *FP*, and *FN* indicate number of true positives, false positives, and true negatives, respectively. Since biological networks are often sparse and therefore imbalanced, we also tested the optimal threshold using the Matthews Correlation Coefficient (MCC) [Chicco and Jurman, 2023]:

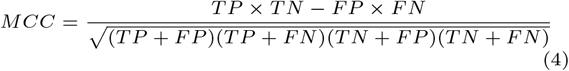

Different metrics capture distinct aspects of the inferred network, and no single evaluation metric is sufficient to fully assess the quality of an NIA. Thus, the evaluation metrics we considered fall into two categories: pairwise interactions and centrality measures.

### Pairwise Interaction Measures

*Area Under the Receiver Operating Characteristic Curve* (AUROC) and *Area Under the Precision-Recall Curve* (AUPR) are evaluation methods that assess performance by scanning across all possible threshold values applied to association matrices, generating curves that balance two desirable properties.

The ROC curve reflects the trade-off between the *True Positive Rate* (*TPR* = *TP/*(*TP* +*FN*)) and the *False Positive Rate* (*FPR* = *FP/*(*TN* + *FP*)). The *TPR* represents the proportion of true positive classifications correctly captured in the predicted network. The *FPR* indicates the proportion of false positives included. Ideally, the *TPR* is 1, and the *FPR* is 0.

AUROC is a widely used metric for evaluating classification performance, especially in diagnostic tests. While effective at distinguishing between ‘bad’ and ‘good’ models, AUROC has been shown to lack sensitivity in differentiating among ‘good’ models [Marzban, 2004]. Additionally, AUROC performs poorly on imbalanced datasets, such as sparse binary networks commonly encountered in metabolomics [Boughorbel et al., 2017].

As a complement to ROC, *Precision-recall* (PR) is widely used for evaluating binary classifiers. PR reflects the trade-off between the *Positive Predictive Value* (PPV, or precision) and the *True Positive Rate* (TPR, or recall). The PR curve is constructed similarly to the ROC curve, replacing TPR and FPR with PPV and TPR, respectively.

AUPR provides a single summary statistic of classifier performance. Given the imbalanced nature of binary biological networks, where non-edges greatly outnumber edges, AUPR is increasingly preferred over AUROC for assessing network inference algorithms in bioinformatics [Saito and Rehmsmeier, 2015].

MCC [Matthews, 1975] is widely used for binary classification, particularly for evaluating network inference methods on imbalanced data, where it often outperforms AUROC [Boughorbel et al., 2017]. MCC quantifies the correlation between reference and predicted classifications, with values ranging from *−*1 to 1 [Chicco and Jurman, 2023]. A value of +1 indicates perfect prediction, *−*1 indicates complete disagreement, and 0 suggests performance equivalent to random classification.

### Centrality Measures

Four centrality measures were selected to summarize the inferred networks, informed by prior work [Liu et al., 2020]. *Degree centrality* measures the number of connections a node has, with higher values typically indicating greater importance in the network. This measure is normalized by the total number of nodes. Many biological networks, including metabolic networks, exhibit a scale-free structure where node degrees follow a power-law distribution. This results in a network with many weakly connected nodes and a few highly connected hubs [Liu et al., 2020].

*Betweenness centrality* assesses the influence of a given node on the flow of communication between other nodes, highlighting nodes that serve as critical intermediaries [Freeman, 1977]. The betweenness of a given node (*v*_*i*_) is defined as:

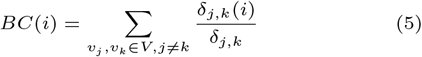

where *V* is the set of network nodes, *δ*_*j*,*k*_ is the number of the shortest paths from nodes *j* to *k* and *δ*_*j*,*k*_(*i*) is the number of these shortest paths that pass through node *v*_*i*_.

*Closeness centrality* measures the importance of a node based on its average distance to all other nodes in the network. Nodes with higher closeness centrality can propagate information more efficiently, requiring fewer hops to reach other nodes. The closeness centrality of a given node *v*_*i*_ can be calculated as the mean value of the inverse of the distance from *v*_*i*_ to other nodes:

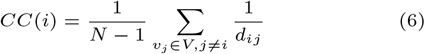

where *N* is the total number of nodes in the network, and the distance between two nodes, *v*_*i*_ and *v*_*j*_, *d*_*ij*_ is defined as the shortest path (minimum number of edges to be traversed) between them [Liu et al., 2020]. If no path between the two nodes exists, the closeness is 0.

*PageRank centrality* (*R*) measures the importance of a node based on the importance of its neighbors. Values are iteratively updated over *t* iterations or until convergence, as defined by the following equation:

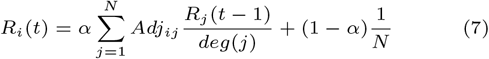

where *α* is the probability of being influenced by a neighboring node, *Adj*_*ij*_ indicates the presence of an edge between nodes *i* and *j*, and *deg*(*j*) is the degree of node *j* [Liu et al., 2020].

PageRank is primarily designed for directed graphs but has been adapted for undirected graphs by treating each undirected edge as bidirectional. PageRank guarantees convergence and avoids degenerate behaviour that can arise in graphs with disconnected components [Liu et al., 2020].

Using centrality values to assess network inference performance offers the advantage of incorporating the overall network structure. However, a key challenge is their sensitivity to errant edges. For instance, a well-reconstructed network with a single additional edge between distant metabolites can significantly increase closeness values compared to the ground truth.

Here, we leveraged the fact that each inferred network shares the same nodes, allowing us to score connectivity on a node-wise basis against centrality measures derived from the ground truth. Centrality measures were calculated for each node in the reference network and compared to those from the inferred network. Differences were summarized by mean absolute error (MAE) and repeated 100 times to show variance. A perfectly recovered network will have a residual error of 0.

### Aspirational Network

To better understand the nuances of evaluation methods across varying signal levels, we introduce here an *aspirational network*, which allows precise control over the amount of noise obscuring the true signal. This approach isolates the effect of sample size by providing the true adjacency matrix with added noise, eliminating the influence of biological outliers and NIA-specific limitations. While there is no exact formula to translate sample size into signal quality, we assume signal quality improves with an increasing number of samples.

We define the aspirational network with a signal level ranging from 0 to 1. At all signal levels, the association matrix was initialized as a uniformly distributed random matrix with values between 0 and 1 (*U*_[0, 1]_). The signal level adjusts the matrix by incorporating a proportion of the ground-truth interacting edges:

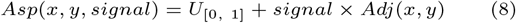

where *Adj*(*x, y*) is 1 if an interaction exists between metabolites *x* and *y* in the reference adjacency matrix, and 0 otherwise. The resulting association matrix values is strictly positive and range from 0 to 2. The aspirational network was generated using only the adjacency matrix from the AA model as input.

### Network Inference Algorithms Evaluated and Their Assumptions

#### Correlation-based Methods

Correlation-based methods are widely used in network inference due to their computational efficiency and scalability for large datasets [Saint-Antoine and Singh, 2020]. These methods quantify pairwise associations between metabolites, offering a simple and intuitive approach to constructing networks. However, a key limitation is their inability to distinguish between direct and indirect interactions, often leading to system-wide correlations that obscure the true network structure [Krumsiek et al., 2011].

**Pearson’s correlation** (*CORR*_*P*_) is one of the most common correlation-based methods. It assumes a normal data distribution and identifies linear relationships between variables [Pearson, 1920]. The correlation coefficient ranges from *−*1 to 1, with *CORR*_*P*_ (*x, y*) = *±*1 indicating perfect linear dependence and *CORR*_*P*_ (*x, y*) = 0 indicating complete linear independence. While its simplicity is appealing, Saccenti et al. [2019] highlight that measurement errors bias correlation coefficients toward zero, potentially impacting the significance of inferred edges.

**Spearman’s correlation** (*CORR*_*S*_) can be considered an extension of *CORR*_*P*_ wherein the data is converted to ranks before calculating the coefficient [Spearman, 1904]. Spearman’s correlation makes no assumptions of the underlying data distribution and can detect both monotonic and linear relationships. Similar to Pearson’s correlation coefficient, a value of *CORR*_*S*_(*x, y*) = *±*1 indicates a perfect monotonic relationship between *x* and *y* while *CORR*_*S*_(*x, y*) = 0 indicates monotonic independence. Since biological systems are governed by non-linear interactions, Spearman correlation is preferred given sufficient sample size [Camacho et al., 2005].

**Kendall’s correlation** (*CORR*_*K*_) is similar to Spearman’s correlation, in that it measures the strength of associations based on the relative ordering of data [Kendall, 1938]. Kendall’s correlation is less sensitive to outliers and provides more reliable *p*-values with smaller sample sizes, making it advantageous in these contexts.

**Biweight midcorrelation** (*bicor*) is designed to be more robust to outliers compared to *CORR*_*P*_, which is particularly valuable when working with small sample sizes. The *bicor* method has been successfully applied in areas such as gene regulatory network inference and differential coexpression analysis [Zheng et al., 2015]. Similar to other correlation measures, *bicor*(*x, y*) produces coefficients ranging from *±*1, with 0 indicating independence.

**Partial correlation** (*pcor*) quantifies the relationship between two variables after accounting for their linear dependencies on other variables, making it useful for distinguishing direct from indirect associations. Partial correlation has been used in metabolic network inference but are computationally expensive and require a larger number of samples [Krumsiek et al., 2011].

Previous studies have provided key insights into the expected performance of correlation methods in different situations. *CORR*_*S*_ performs well in metabolomic studies due to its reliance on monotonicity rather than linearity given a sufficient number of samples. When sample sizes are limited, it has been recommended to use *CORR*_*P*_ with a high significance threshold [Camacho et al., 2005].

Moreover, Santos et al. [2014] have evaluated the strengths of various correlation methods using both simulated and experimental gene expression data. Their findings suggest that when monotonicity can be assumed, *CORR*_*S*_ or *CORR*_*K*_ are preferable to *CORR*_*P*_, as they can detect both linear and non-linear monotonic relationships with greater statistical power.

#### Information Theoretic-based Methods

Information theoretic methods use **mutual information** (*MI*) to find pairwise scores for interactions. Similar to *CORR* methods, *MI* can be used on its own to produce weighted networks or with thresholding to produce binary networks [Butte and Kohane, 1999]. The mutual information *MI*(*x, y*) between two metabolites *x* and *y* can be calculated as [Song et al., 2012]:

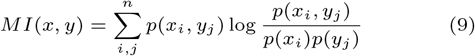

where *p*(*x*_*i*_, *y*_*j*_) is the joint probability distribution function of *x*_*i*_ and *y*_*j*_ and *p*(*x*_*i*_) and *p*(*y*_*j*_) are the probabilities that *x* = *x*_*i*_ and *y* = *y*_*i*_, respectively. Entries of an association matrix generated using *MI* can have values in the range of [0, +*∞*]. *MI* presents a theoretical advantage over correlation-based approaches by capturing non-linear relationships, which are known to be present in biological systems [Camacho et al., 2005]. However, it is important to note that recovering non-linear relationships with limited samples is challenging, and estimating *MI* from small sample sizes introduces significant errors [Song et al., 2012].

The **Context Likelihood of Relatedness** (*CLR*) algorithm was designed as an extension of *MI* [Faith et al., 2007]. The *CLR* algorithm estimates the likelihood of the *MI* score for a given pair of metabolites against the background of the *MI* scores for that pair, minimizing the effect of indirect interactions. *CLR* is often used as a post-processing step for any association matrix.

**MRNET** evaluates an *MI* association matrix for inference based on the maximum relevance/minimum redundancy (MRMR) algorithm [Ding and Peng, 2005, Meyer et al., 2007]. The MRMR algorithm iteratively selects a feature that has maximum relevance with respect to the target variable and minimum redundancy relative to the features that have been selected in previous iterations [Ding and Peng, 2005]. As a feature selection technique, MRMR has been successfully applied to disease biomarker identification [Mahendran et al., 2021]. In its original publication, *MRNET* was shown to perform competitively against *CLR* on 30 simulated microarray data sets [Meyer et al., 2007].

#### Regression-based Methods

Regression-based methods quantify relationships between metabolites by solving regression problems to predict a target metabolite based on others. These methods can infer potential causal relationships [Saint-Antoine and Singh, 2020] and several have been benchmarked on DREAM5 [Marbach et al., 2012].

**Gene Network Inference with Ensemble of Trees** (GENIE3) uses variable selection with ensembles of regression trees to infer associations. Its tree-based approach allows *GENIE*3 to detect non-linear relationships [Huynh-Thu et al., 2010]. Briefly, biological samples are represented as vectors, with features corresponding to metabolites. To determine associations for a target metabolite *x, x* is masked, and regression trees are trained to predict it based on the remaining metabolites. Metabolites most important for predicting the masked values are identified as being highly associated with *x*. The ensemble of trees ranks the importance of each input metabolite. For consistency with other methods used in this work, directionality was removed by taking the maximum of *A*_*x*,*y*_ and *A*_*y*,*x*_.

#### Resampling-based Methods

The **Probabilistic Context Likelihood of Relatedness of Correlation** (*PCLRC*) algorithm extends *CLR* by iteratively resampling subsets of the available data. In each iteration, the top *Q*% of highest-associated metabolite interactions are recorded. The final association matrix reflects the fraction of times each metabolite pair appeared in the top *Q*% interactions. Following the original procedures [Saccenti et al., 2015, Jahagirdar et al., 2019], each resampling used 75% of the available data, and the top 30% of interactions were saved. Although 10^4^ repetitions were originally recommended, here we limited the number to 10^3^ due to the computational demands of applying *PCLRC* across multiple association matrices.

### Computational Settings

Five hundred synthetic samples were generated from the simulated AA model using MATLAB (R2022a). Each NIA utilized subsets of these samples with sizes *n* = [5, 10, 20, 50, 100, 200]. For each method and sample size, 100 repetitions were conducted. We included subsets of size 100 and 200 to evaluate the NIAs when the number of samples was greater than the number of metabolites.

We followed the approach laid out in Jahagirdar et al. [2019] to ensure we reached steady state by running the simulation for 90 hours. As with Jahagirdar et al., variation between profiles was achieved by slightly perturbing the reaction parameters via uniformly resampling from *±*10% of the parameters’ optimized values. Association matrices were generated for each algorithm from the following sources: SciPy [Virtanen et al., 2019] provided Kendall-Tau and Spearman correlations. The Python version of GENIE3 [Huynh-Thu et al., 2010] was obtained from Github. Numpy had the most optimized Pearson correlation algorithm in terms of wall-clock time and core utilization [Harris et al., 2020]. Biweight midcorrelation was rewritten from Pingouin to speed up the algorithm by compilation [Vallat, 2018]. MRNET was rewritten due to challenges with implementing the official source in this workflow. *CLR* and *PCLRC* are processes applied to an association matrix, and were rewritten to allow flexibility in this workflow. Partial Correlation was performed with the Python package Pandas [McKinney, 2010]. Sci-kit Learn [Pedregosa et al., 2011] was used for calculating *MI* by way of entropy with k-nearest neighbours, where *k* defaults to 3. The rewritten algorithms were accelerated using Numba compilation [Lam et al., 2015].

### Simulated Sarcopenia

To test whether inferred networks could be used to differentiate biological conditions, we adapted the AA model to examine the lipidomics of sarcopenia, an age-dependent skeletal muscle wasting syndrome. We informed our AA model using the data and conclusions of Palla et al. [2020] comparing capacity of different correlation-and regression-based network approaches to infer the underlying sarcopenia-associated enzymatic change revealed by these data. Palla et al. used high-performance liquid chromatography tandem mass spectrometry (LC-MS/MS) to measure prostaglandin family members in young and aged murine skeletal muscle and then identified the causal enzymatic expression changes responsible for measured age-dependent differences in prostaglandin abundances. They found that an increase in 15-hydroxyprostaglandin dehydrogenase (15-PGDH) in aged tissue contributed to skeletal muscle atrophy. The 15-PGDH enzyme catalyzes two reactions in our network: Prostaglandin E2 (PGE2) → 15-keto-PGE2, and prostaglandin D2 (PGD2) → 15-keto-PGD2. They demonstrated that a reduction in PGE2 and PGD2 levels in aged tissue was due to an increase in their degradation to 15-keto-PGE2 and 15-keto-PGD2 respectively. The authors further demonstrated that it was the resulting loss of PGE2-associated signalling that, in part, caused sarcopenia-associated muscle wasting.

To simulate the young and aged states in prostaglandin abundances in our network, we started from the reaction parameter values reported by Jahagirdar et al. [2019] and amplified *V*_*max*_ by 1000 times, informed both by the increase in 15-PGDH mRNA observed by Palla et al. [2020] and prior work correlating mRNA to protein levels [Lahtvee et al., 2017, Buccitelli and Selbach, 2020]. *K*_*m*_ was also increased by 2.5 times for the two affected reactions. We generated 200 samples from these simulated young and aged networks to obtain the largest sample sizes considered in this study. Each sample was initialized from a steady-state. Enzyme parameters were updated using the *±*10% method, then simulated for 20 hours to reach their new equilibrium using COPASI [Hoops et al., 2006].

### Bootstrapped Network Differentiation

To find differences between the aged and young inferred networks, we took inspiration from *PCLRC* to minimize the impact of sample selection. By generating many possible networks from a subset of the available data, we identified edges that occurred reliably (i.e., in *>* 50% of resampled networks). We used these reliable edges to define a candidate network, that was less impacted by sampling bias. Next, to create candidate networks for sarcopenia-simulated datsets, we started with 200 samples from the young distribution. One hundred samples with replacement were selected to infer intermediate networks.

These intermediate networks were subsequently separated into two groups. For each group, the candidate network was summarized as the edges that occurred in *>* 50% of the intermediate networks. By comparing the two groups from the young distribution, we observed the degree of variation between simulations. This same process was repeated to generate an aged distribution.

## Results

### Performance Measure Traits

With the performance measures defined and exploiting the aspirational network (see Section Aspirational Network) to control signal levels, we investigated how performance approximated function. Fifty signal levels between 0 and 1 were selected with increments of 0.02. For each signal level 100 aspirational association matrices were generated.

Consistent with previous work demonstrating that AUROC does not utilize the full metric range (0-1) [Marzban, 2004], we found that values saturated early (i.e., at relatively low signal level; Supplementary Figure 1A), indicating AUROC was unable to differentiate between multiple ‘good’ networks. Our data also confirmed that AUPR values were saturated at both signal-to-noise extremes (Supplementary Figure 1B). Taken together, both area under curve metrics showed low variance across signal-to-noise levels (Supplementary Figure 1A,B).

FDR has high variances when signal is low, though this appears to be resolved at even very low sample sizes (Supplementary Figure 1C). MCC has been shown to perform well for low signal-to-noise networks, did not contain discontinuities, and had nearly uniform variance throughout the signal range (see Section Evaluation Metrics). Here, there was a slight saturation of the measure at high signal (Supplementary Figure 1D). Using FDR and MCC, we also compared the impact of optimizing the threshold for F1-score or MCC score. We found that the difference was minimal. As such, going forward, we used the F1-score for optimizing the threshold. Visualizations of the pairwise evaluation measures are available in the Supplementary Material (Supplemental Figure 1).

Centrality evaluation measures were also inspected for their behaviour using the aspiration network algorithm with varying signal levels (Supplemental Figure 2). Betweenness centrality had a discontinuity in the curve, likely resulting from biologically incorrect edges bringing metabolites together that should be apart in the low signal regime. Properly recovered edges began to appear at higher signals (*>* 0.6). Closeness centrality showed a stark discontinuity at low signal that we again attributed to random edges confusing this metric. While degree centrality starts with high variance, it quickly converged to a low error with signal *>* 0.2, making it potentially useful for differentiating NIAs during evaluation and development. The PageRank performed well across the signal ranges *>* 0.2, with a nearly linear relationship to signal.

**Fig. 2.**
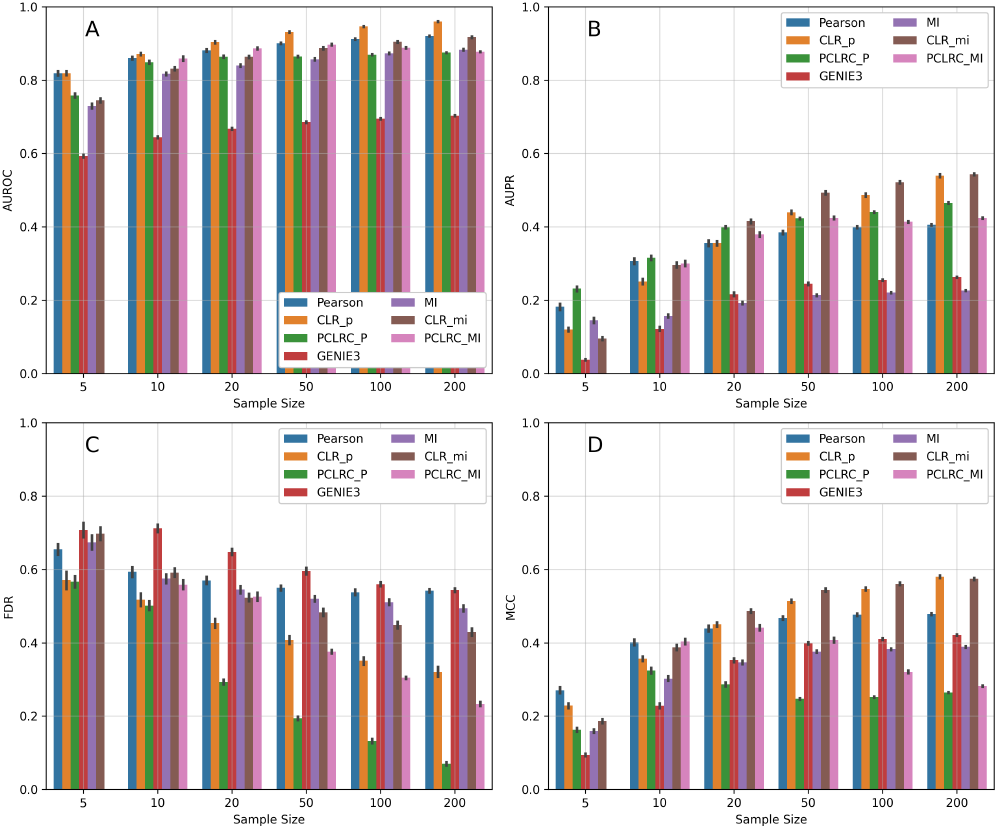
Pairwise metric evaluations for a selection of Network Inference Algorithms. A) AUROC. B) AUPR. For C) FDR and D) MCC a threshold was selected that optimized for F1-score to ensure the best possible result for cross-evaluation.

### Network Inference Evaluation

The previous section provided insight into how various performance measures were affected by noise. It is commonly assumed that additional samples will reduce noise and thus reveal the signal required to infer a metabolic network [Suarez-Diez and Saccenti, 2015]. We show here that this assumption is not relevant for reasonable experimental sample sizes, and that even large sample sizes do not converge to reveal the underlying network structure.

### Pairwise Measures

In addition to comparing performance measure traits, Figure 2 shows a subset of NIAs for visual clarity. The full set of NIAs tested are available in Supplementary Materials. *PCLRC*_*MI*_ was omitted from the 5 sample size evaluations due to *MI* requiring at least 3 samples and *PCLRC* selecting a subset of those available. As discussed above, AUROC (Figure 2A) began to converge for higher sample sizes at approximately 0.9 for most methods. Knowing that the AUROC measure saturates at 1 with signal levels as low as 0.6, the convergence to a lower value indicated systematic differences in the inferred network relative to the reference network. We saw an increase in AUPR as the number of samples increases, which is to be expected (Figure 2B). Post-processing methods (*PCLRC* and *CLR*) typically achieved higher AUPR over the raw-association methods. The highest AUPR achieved was from *CLR* methods at 200 samples, though no method reached an AUPR of *>* 0.6 (Figure 2B). With additional samples, FDR (Figure 2C) decreased incorrect edge detection and post-processing NIAs typically accelerated this trend. The low variance of FDR at these very small sample sizes suggested that the high variance at low signal is an artifact that can be ignored. *PCLRC* methods are much better able to remove outlier influence with larger sample sizes than the corresponding raw method (*≥* 50 samples); however, these approaches provide only marginal improvement with low sample sizes (*≥* 10) (Figure 2D). For NIAs that use the raw association output, FDR plateaued at 0.5. Finally, using MCC, the raw output NIA models showed a plateau at 0.5 that was approached early and was not overcome (Figure 2D). *CORR*_*P*_ performed best at low sample sizes (*≤* 10). *PCLRC* methods peaked at sample sizes around 20 and decreased with more samples. This surprising result was further explored in Supplementary Materials: PCLRC Algorithmic Limit. We show that this early performance peak appeared to be caused by saving only the top interactions for each resampling. NIAs using *CLR* were the only methods that crossed 0.5 for this metric, and only when the number of samples was very high (*≥* 50, Figure 2D). As MCC is designed to avoid the challenges of imbalanced data, failing to perform well here shows that none of these NIAs appreciate the sparse connectivity of the underlying network.

### Centrality Analysis

The true network adjacency matrix was used to calculate reference centrality scores for each of the four centrality methods providing 4 vectors with 83 elements. A selection of NIAs were evaluated on these centrality measures: *Pearson* and *MI* representing raw algorithm output, and *CLR* and *PCLRC* representing post-processing methods. Figure 3 visualizes the results repeated 100 times for each NIA method at each sample size.

**Fig. 3.**
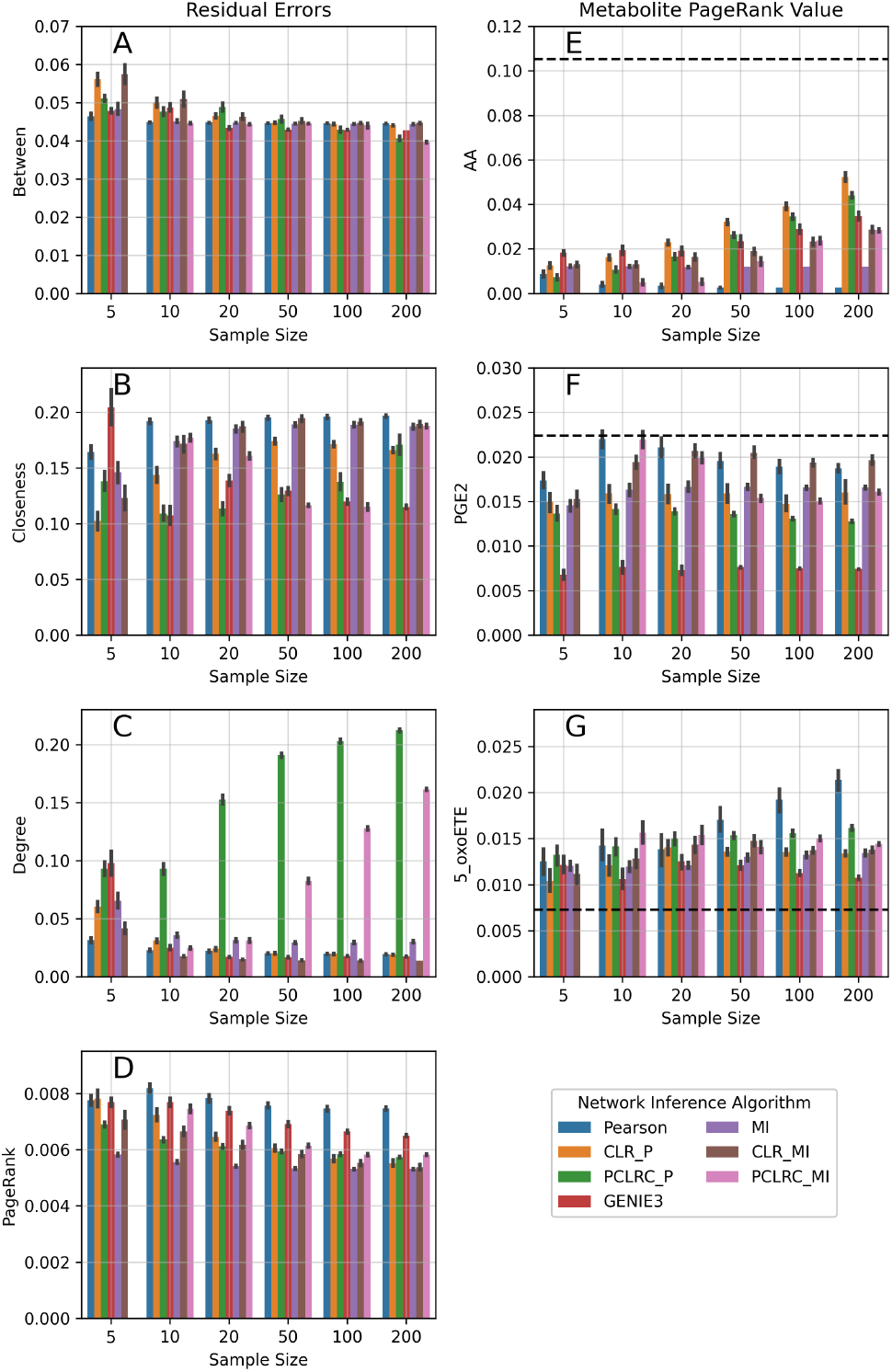
Centrality metrics for a selection of NIAs. Panels A-D depict the differences between the centrality vector from inferred networks and the reference network. Zero indicates the inferred network matches the reference network. Panels E-G show the PageRank centrality for individual metabolites: AA (core metabolites), PGE2 (intermediate in the network), and 5-oxoETE (distal metabolites). The black line indicates the PageRank value of these metabolites in the reference network.

*Betweenness* showed marginal improvement with additional samples regardless of NIA method (Figure 3A). *PCLRC* methods showed a drop at 200 samples, but the overall trend converged to a different point from the reference network. *Closeness* (Figure 3B) had high variance between runs, and many methods appeared to be diverging from the reference network as the number of samples increased. *Degree* centrality (Figure 3C) indicated that *PCLRC* methods diverge with additional samples. *PageRank* showed limited improvement with additional samples (Figure 3D). *MI* had the lowest PageRank error at 200 samples, but the error was comparable 5-200 samples indicating that increasing sample size did not improve outcome.

### Metabolite Inspection

We next assessed where errors accumulated across the networks. We isolated three metabolites at different points in the reference network: AA representing a central node, PGE2 representing an intermediate node, and 5-oxoETE representing a distal node. Figure 3E-G separates these metabolites by sample size with the dotted black line indicating the PageRank value of that node in the reference network. The PageRank of the corresponding metabolite was calculated across 100 repetitions for each of the tested NIAs. AA (Figure 3E) showed significant underestimation of the importance of this metabolite regardless of NIA. PGE2 importance (Figure 3F) was rarely correctly estimated, although it is important to note the high error bars and failure for any method to converge with more sample sizes. Finally, 5-oxoETE (Figure 3G) was overestimated by all methods. Collectively, these results suggested a flattening of PageRank values across the inferred networks. This flattening of node importance on the inferred network was visualized in Figure 4. Here, the AA network was inferred using *CLR*_*P*_ with 200 samples. Nodes were placed in the same positions as the reference network (Figure 1), and coloured by the error in PageRank estimation. This representation clearly demonstrates that nodes towards the exterior of the network were systematically overestimated while highly central nodes were underestimated.

**Fig. 4.**
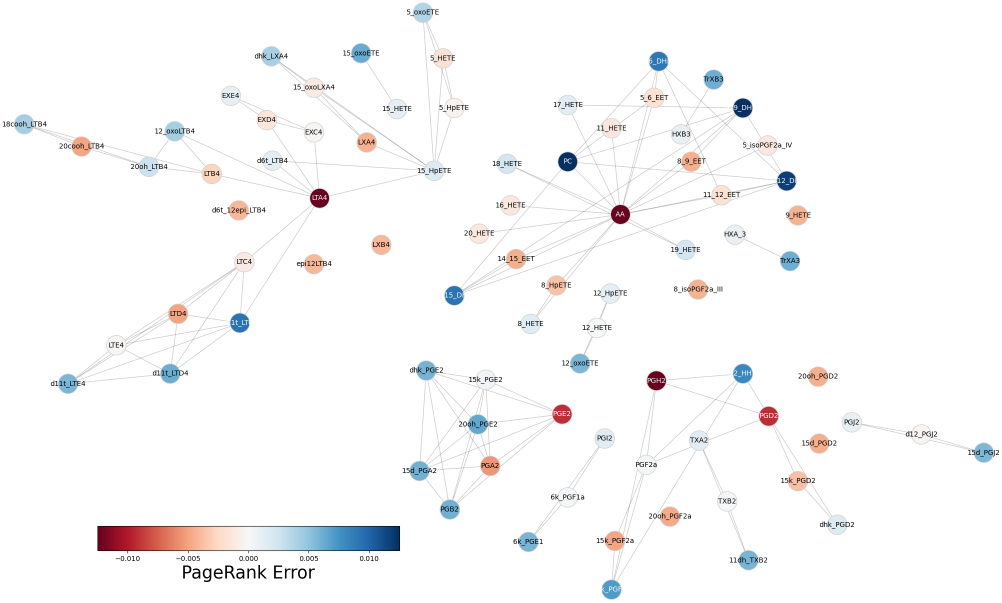
A single example of a recovered network using *CLR*_*P*_ inference with 200 samples. Nodes are coloured by their disagreement with the PageRank value calculated for the reference network, with red indicating underestimated values and blue indicating overestimated values.

### Bootstrapped Networks

To explore the interactions reliably predicted by NIAs we adapted the bootstrapping methodology of *PCLRC*. By inferring networks from a subset of the data and selecting the edges that occurred in the majority of networks, we limited the impact of sampling variation. See section *Bootstrapped Network Differentiation* for more detail on the methodology.

We found that candidate state networks were internally consistent by creating two candidates states from 50 repetitions and investigating the difference in inferred connections (Supplementary Figure 5A,B). We found 4 edges were in disagreement and 112 edges in agreement in the young network. Five edges were in disagreement and 127 in agreement in the aged network. The interactions between young and aged showed 38 differentially inferred edges (Supplementary Figure 5C). Figure 5 visualizes these edges disagreement and identifies distributional change as the source of these disagreements. Edges marked in blue indicate edges that were found in the young but not aged distribution. Red marks indicate edges present in aged but not young distributions. These visualizations clearly show that the edges directly affected by the 15-PGDH enzyme (black dotted lines) were not identified by this analysis because these reactions were inferred in both distributions. In fact, the distributional shift of inferred edges between the two states included changes in edges far from the 15-PGDH enzyme.

**Fig. 5.**
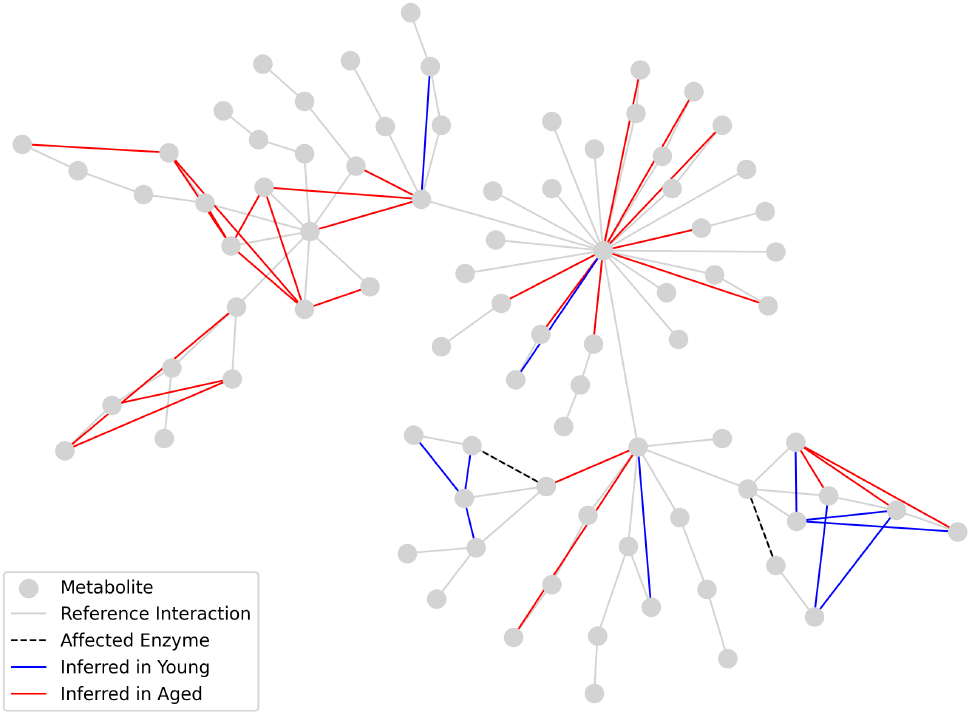
Edges that are in disagreement between the young and aged distributions. Gray lines depict the interactions defined by the reference network between metabolites (gray nodes). Black dotted lines show the two 15-PGDH enzyme reactions. Blue lines indicate edges that were regularly found in the young network, but not the aged network, whereas red lines indicate the opposite. Note that the *bona fide* enzymatic disruption identified by Palla et al. [2020] was not identified as a disagreement between the young and aged networks.

## Discussion

Using a simulated metabolic network of AA degradation, we demonstrated that NIAs, without additional network knowledge or constraints, do not recover the underlying metabolic network. We expanded this assessment using a ‘real-world’ metabolic disruption, age-dependent sarcopenia, demonstrating that NIAs were unable to highlight the primary metabolic disruption.

We started by characterizing the limits of network evaluation measures by introducing an aspirational network that allowed us to control the total noise. The output of any NIA is limited by the number of samples (analogous to the fraction of adjacency matrix signal), and therefore ‘aspires’ to the aspirational network given the assumption that the NIA will converge to the underlying metabolic network. Our aspirational network is parallel to Diez et al. [2015] who looked at the number of samples required for network convergence. They, however, were interested in convergence to the inferred network when the number of samples was not a limiting factor, instead of converging to the underlying network, which they did not have access to.

We tested several popular NIAs that find pairwise associations between metabolite concentrations to infer the underlying network using varying sample sizes. Determining the quality of the inferred networks relied on both edge-level and network-level performance measures. We found that while the edge-level performance measures often improved with additional samples (AUPR, AUROC, FDR) there were also algorithmic limitations (MCC; see Supplementary PCLRC Algorithmic Limit). Additional samples caused the evaluation metrics to converge, but not to the expected optimal values of the metric. These performance measure evaluations were under the optimal threshold value, requiring knowledge of the ground-truth interaction network. We chose this idealized methodology to focus on the best-case scenario for each NIA method.

The centrality-based performance measures demonstrated NIAs do not converge to the underlying network more directly; additional samples, even by almost 2 orders of magnitude, had little effect on the simulated case. The surprising stability of centrality measures, regardless of sample size, brought us to interrogate where NIAs were introducing errors. We found PageRank values for central metabolites were systematically underestimated and distal metabolites were overestimated. PageRank was originally designed for directed graphs. It is important to note that while the underlying network is directed, the inferred networks are not. Thus, we compared degree, betweenness, closeness, and PageRank centralities, although we recognized that other approaches could have be applied [Liu et al., 2020].

While our work has not identified what biological relationships NIAs exactly capture, it does add *in-silico* examples to the increasing number of calls that a better understanding of the underlying utility of network inference is needed [Nyamundanda et al., 2013]. In the cases where the underlying metabolic network is known, alternative hybrid methods have shown interesting results [Li et al., 2024]; however, this type of information is not always available.

We do show that, although inferred networks do not recover the underlying metabolic interactions well, they can be used to differentiate disease states (but not necessarily the underlying mechanisms resulting in these stats) as we found that basic shunting of organic mass affects associated connections in the inferred networks. Our results suggest that network inference can be used to differentiate sample distributions, identifying a change of state and/or distinguishable conditions, but that caution should be applied when inferring candidate reactions responsible for this change of state since the impacted reactions can be dispersed in the network, far from the causative reaction/mutation/expression. This utility (and deficit) was clearly demonstrated by simulating age-associated changes in lipid metabolism in sarcopenia informed by experimental results of a previous study [Palla et al., 2020]. A change of state between young and aged networks was identified but the empirically validated causal reaction could not be inferred. Taken together, methods for network differentiation from inferred networks require additional empirical follow-up exploration that, in turn, will provide insights into the nature of NIAs output. It is important to note that network inference methods that incorporate prior knowledge about the underlying network [Li et al., 2024] or apply assumptions about network structure (e.g., sparsity [Lingjærde et al., 2020]) are not benchmarked here. While benchmarking knowledge-informed networks is challenging as the type of included information may not be directly comparable to other methods, such work would be valuable

In summary, this work demonstrates that relying on pairwise NIAs without inclusion of any prior knowledge does not recapitulate the underlying network but can provide evidence that the two conditions are indeed distinct.

## Author contributions statement

FA ran experiments and contributed to writing. HG provided literature review and initial experiments. US assisted in initial experimental design and copy editing. SB, MCC, and HJ conceptualized the work, assisted in experiment design and writing.

## Acknowledgment

We thank and acknowledge the Computational Biology Research and Analytics Lab for invaluable feedback. This work was partially supported by funding from NSERC Discovery Grant (SB, HJ), NRC Digital Health Cluster (MCC, HJ), NRC AI4D Challenge program (SB, MCC).

## Supplementary Materials

### Performance Measure Characterization

We characterized how these evaluation metrics performed under an ideal scenario to account for the additional challenges of sample size error and NIA biases. We defined a ‘best-case scenario’ inference algorithm in Section *Aspirational Network* and we show here the pairwise evaluations in Supplementary Figure 1. For FDR and MCC, each association matrix had a threshold selected to optimize F1-score to create a binarized network. We also used using MCC to determine an optimal threshold and found minimal disagreement when converging to the underlying network (i.e., above signal of *≥* 0.5).

At higher signals, we found AUROC was saturated (approximately zero slope) inhibiting its ability to distinguish networks. AUPR and MCC both showed small variance across the levels of noise and limited saturation at both extremes. The high variability of FDR at low signal appeared not to be an issue even with very small sample sizes, as evidenced by the low iteration error bars in Figure 2 at sample size of 5. MCC was approximately linear with low variance across signal levels and some saturation at the highest levels (i.e., above *>* 0.9).

The aspirational network was also used to evaluate the impact of noise on the proposed centrality based measures. An ideal response to increasing signal would be a linear decrease in error. Supplementary Figure 2 shows the four centrality measures with increasing signal. Betweenness (Supplementary Figure 2A) had a discontinuity in response at relatively high signal strength (*>* 0.6) suggesting susceptibility to noise. Closeness (Supplementary Figure 2B) had an approximately flat response to increasing signal between 0.2 through 0.5 indicating it would be unable to differentiate noisy networks. Degree (Supplementary Figure 2C) had two modes; high error and high variance below signal 0.2, and approximately linear with low variance above this point. Empirically, the high error region is relevant when using a sample size of 5 (Figure 3C). PageRank (Supplementary Figure 2D) was considered to work well due to its nearly linear response beginning at a signal factor of 0.2. We also considered using eigenvector centrality, but found convergence issues due to disconnected subgraphs (data not shown).

### PCLRC Algorithmic Limit

The probabalistic approach of *PCLRC* was hypothesized to perform well at very low sample sizes due to resampling reducing the influence of outliers and we observed this for FDR in Figure 2, exemplified at sample sizes greater than 10. However, in the same figure we found that *PCLRC* peaks for MCC with a sample size of 20, plateaued at a lower value with additional samples.

As defined in Saccenti et al [2015], *PCLRC* is provided a set of samples from which to resample (with replacement). Each subset is used to infer a network and the top k% edges are kept. As the number of samples increases, the network inferred by *PCLRC* converges. When the sample size is high, noise plays a smaller role and the strongest edges can be reliably kept. This causes weak but still meaningful edges to be discarded lowering the overall edge quality. When the sample size is low, high confidence edges can be found, and through the impact of noise, low edges can be observed. We show here that the peak at sample size 20 represented the balance between adding more samples for a better estimate of the underlying network and the overconfidence of *PCLRC* in detecting important edges. To decouple these competing effects, we repeated *PCLRC* methods allowing resampling from the full 500 samples. By resampling from the full set, we reduced the impact of the initial selection caused by sample size.

Supplementary Figure 3 shows evaluation metrics of our ‘unlimited’ *PCLRC*. Both the AUPR and AUROC plateau demonstrated that these performance measures are less impacted by small sample sizes. While consistent performance measure across samples sizes is desirable, neither reached a value indicating high quality network inference. FDR decreased as sample size increased. Unlike AUPR and AUROC, where sample size made little difference, the error in FDR appeared to be caused by estimating using an insufficient number of samples. *PCLRC*_*P*_ showed a significantly faster decrease in FDR than the other methods, and *PCLRC*_*MI*_ showed a significantly slower decrease. MCC presented a decrease in quality with additional samples, following our explanation above.

### Network Differentiation

In the context of network differentiation, Supplementary Figure 4 shows inferred networks from two different metabolomic states. Using 200 samples for each of two states, networks were inferred using *CLR*_*P*_. These inferred networks were overlaid on the reference network. For visual clarity, the names of metabolites are omitted. Both states resulted in disconnected graphs and both reliably identified the modified 15-PGDH enzyme interaction (the edge between the pairs of green nodes). However, the presence or absence of an edge cannot reliably be attributed to state difference or sampling noise, leading us to propose the bootstrap network method in section *Bootstrapped Networks*. This proposed bootstrap method for network differentiation relied on minimizing the sampling noise by repeatedly resampling subsets of the original data. Supplementary Figure 5 presents intermediate steps culminating in Figure 5. The left panels show how much sampling variation was observed when selecting a subset of 100 samples from the available 200 samples. These subsamples were used to create a single inferred network by selecting interactions that occur in *>* 50% of the inferred networks. The young and aged variations found few interactions that disagreed between resamplings (4 and 5, respectively). Due to the fact that these were undirected, the upper left triangle is used to list the interactions in all cases. The disagreement between the two states ultimately included 38 discordant interactions. The difference between these states include distant impacts that are more clearly visualized in Figure 5.

### Full Resolution Images

Supplementary Figures 9-12 show pairwise and centrality evaluation results on all considered NIA methods. The corresponding figures in the paper show a sample of the NIAs to not distract from our primary message that NIAs fail to meaningfully converge to the underlying network (Figures 2 and 3, respectively).

**Supplementary Figure 1.**
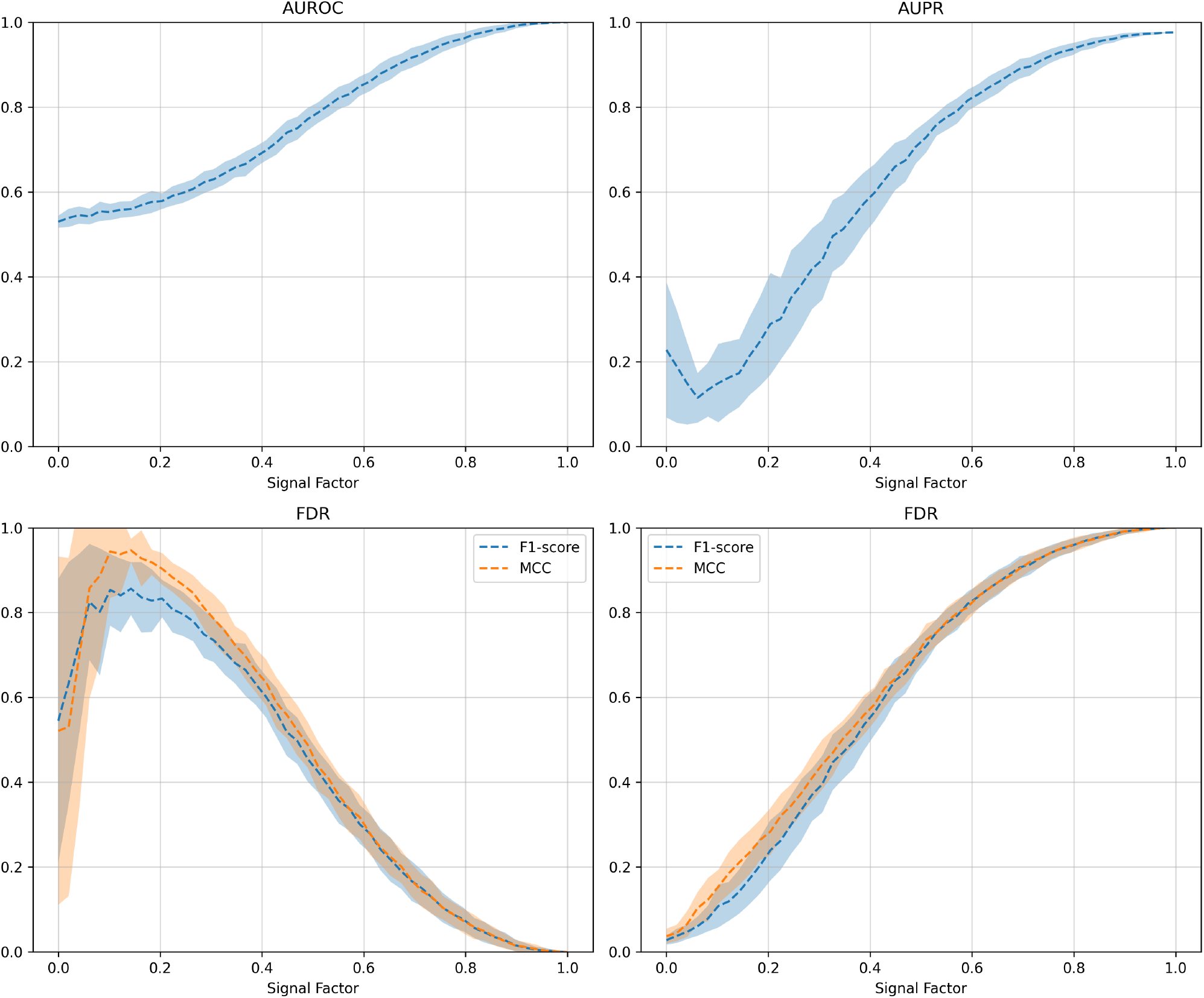
Pairwise metric evaluations of an aspirational network. Optimal threshold values for FDR and MCC were calculated using both F1-score and MCC. Signal levels were repeated 100 times, with dotted line indicating mean value and the shaded region showing one standard deviation.

### Paper Source

The work described in this article were originally explored in the Masters thesis by Haley Greenyer for her degree from the University of Victoria [Greenyer, 2022]. The thesis presents similar evaluation metric testing including the use of graph centrality measures, but does not consider the convergence issues of the network inference algorithms.

**Supplementary Figure 2.**
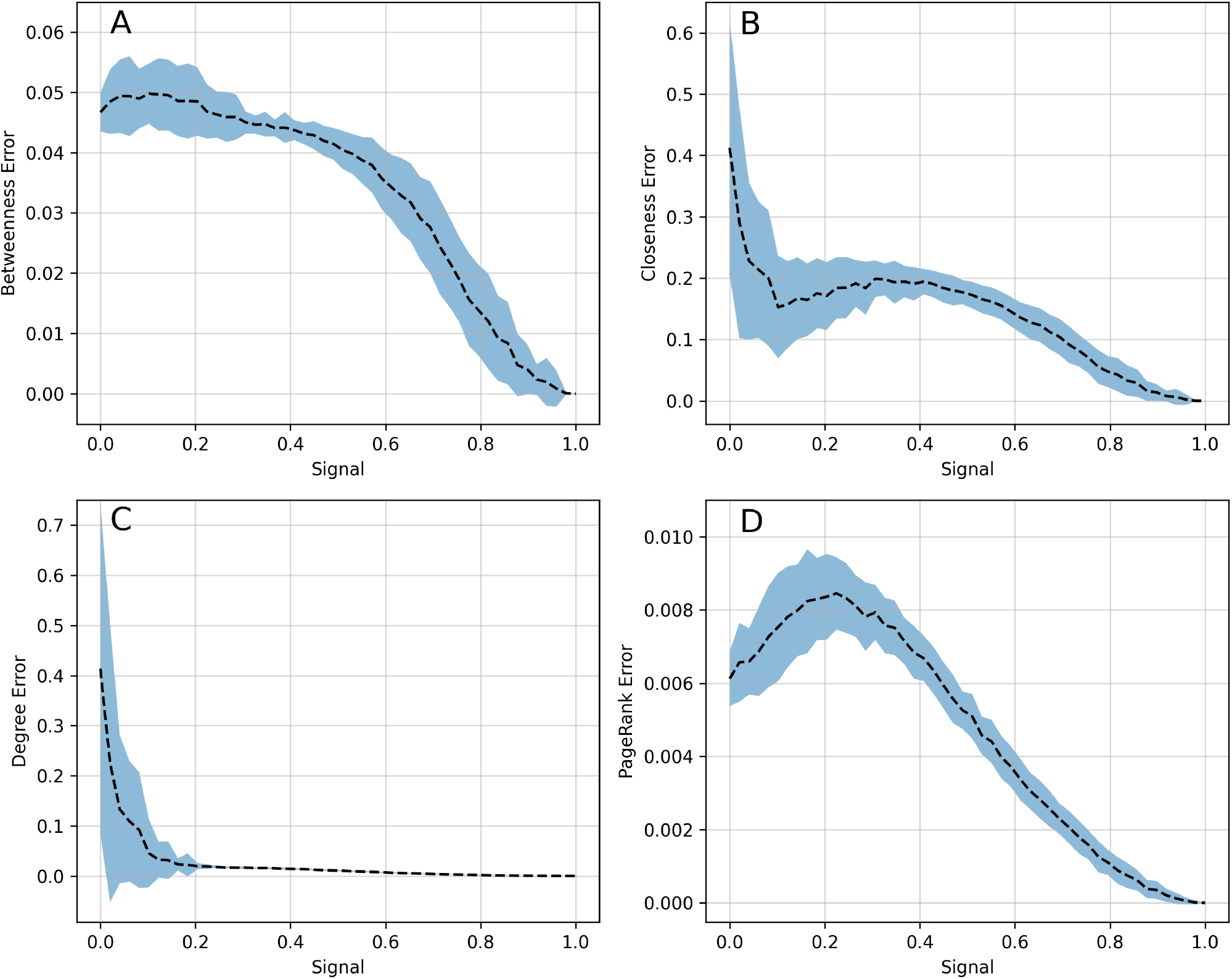
Network Structure evaluations for an aspirational network. Optimal threshold values were calculated using F1-score. Signal levels were repeated 100 times and in increments of 0.02. Dotted lines indicate mean value and the shaded region showing one standard deviation.

**Supplementary Figure 3.**
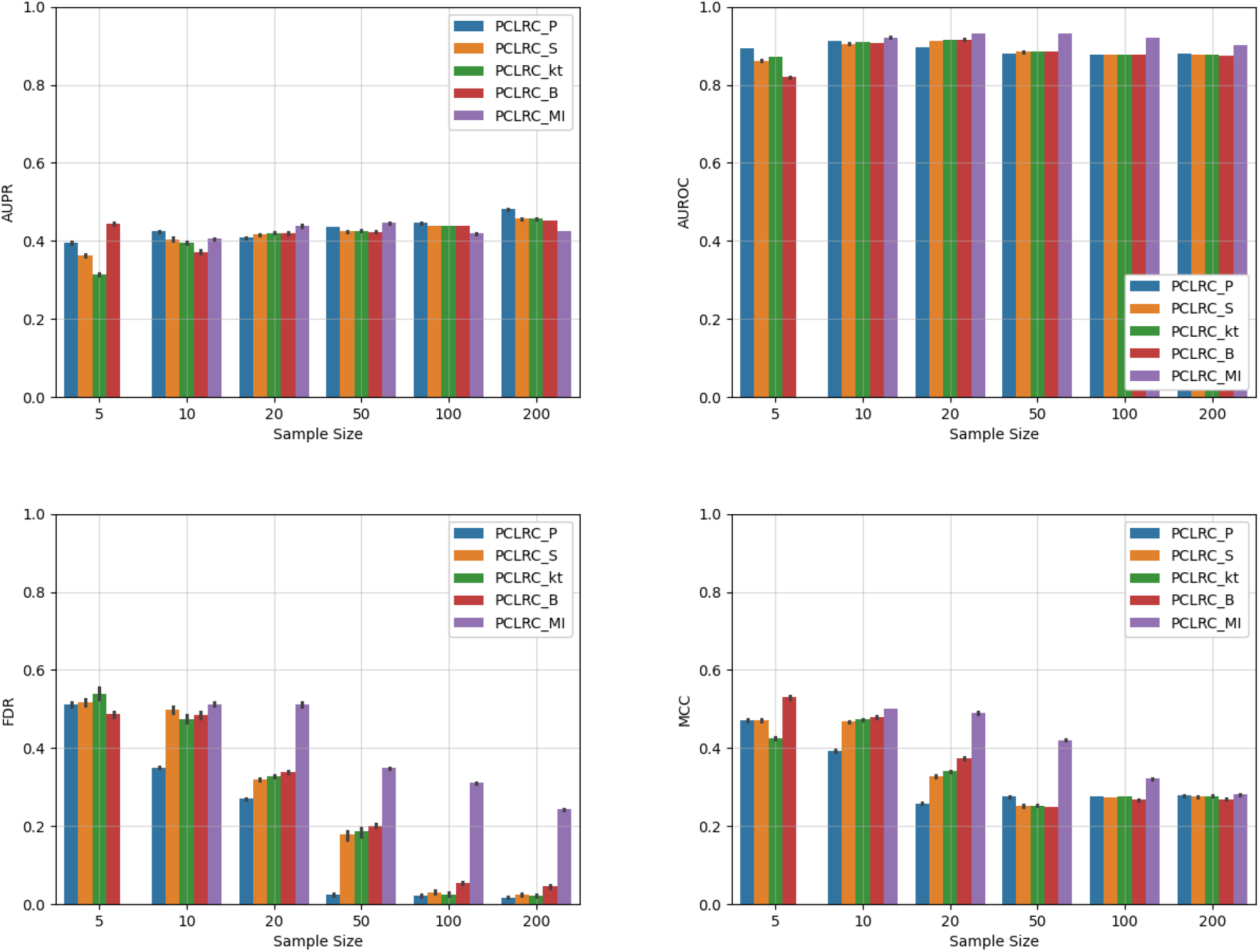
*PCLRC* pair-wise results when resampling from entire dataset. Each method is replicated 100 times at each sample size.

**Supplementary Figure 4.**
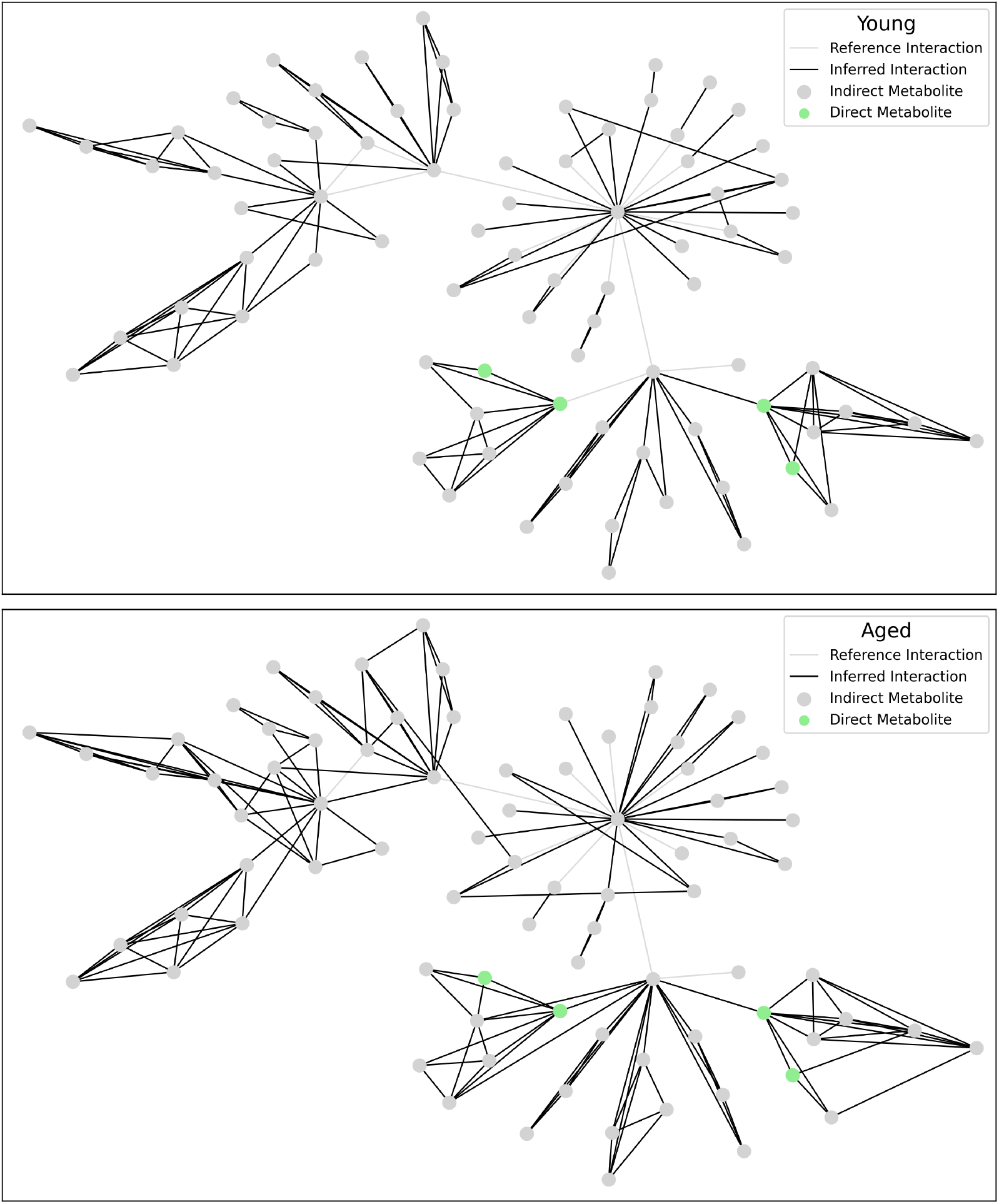
A pair of inferred networks from young and aged datasets. Gray lines show ground-truth edges not found by the inference algorithm. The top graph was inferred from 200 samples from the young distribution of samples; the bottom from the aged distribution. Metabolites directly impacted by the mutation of 15-PGDH are shown in green.

**Supplementary Figure 5.**
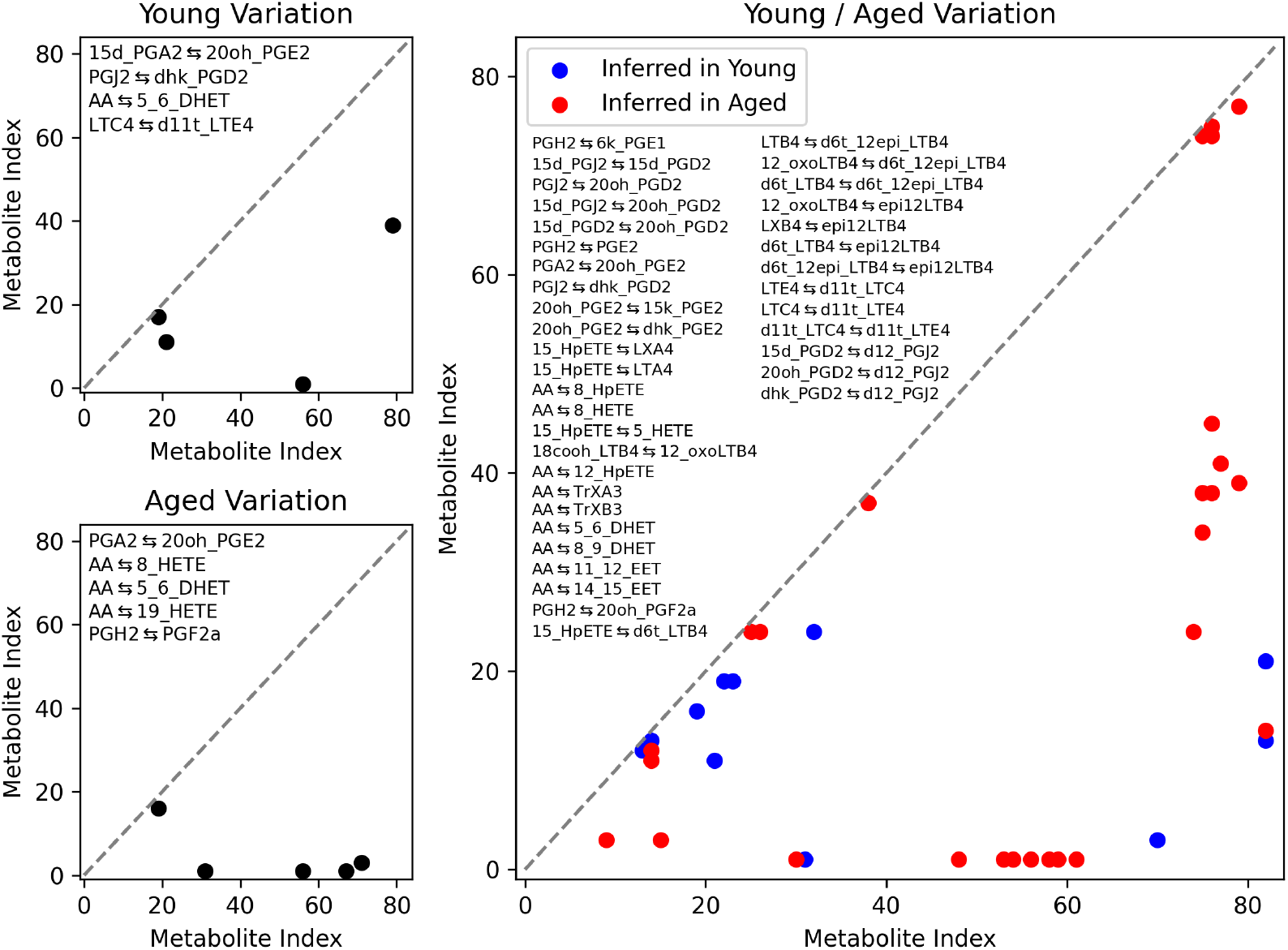
Bootstrapped network differentiation. Young variation shows 4 edges were in disagreement when comparing two bootstrapped networks. Aged variation found 5 edges disagreed. Young/Aged variation shows 38 edges disagree when comparing a young and aged network. Edges that exclusively occur in young are shown in blue; edges exclusively in the aged network are in red. For all panels the reactions in disagreement between networks are written in the upper left.

**Supplementary Figure 6.**
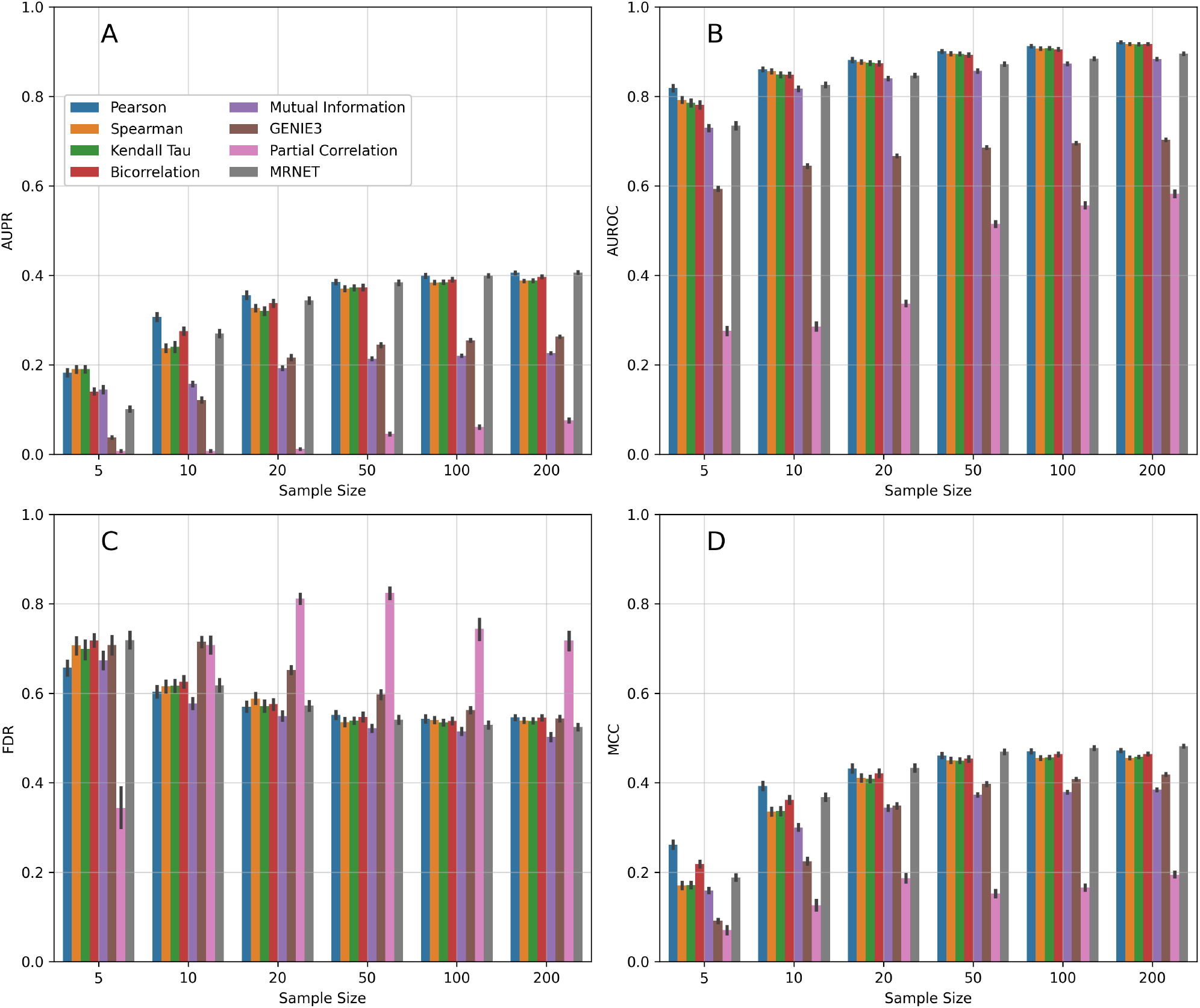
Network inference algorithms (NIAs) evaluated on pairwise performance measures. Thresholding values for FDR and MCC were calculated optimizing for F1-score for each individual network to limit the effect of threshold selection. This selection of NIAs exclude *CLR*-based methods for visual clarity. See Figure 7 for *CLR*-based methods.

**Supplementary Figure 7.**
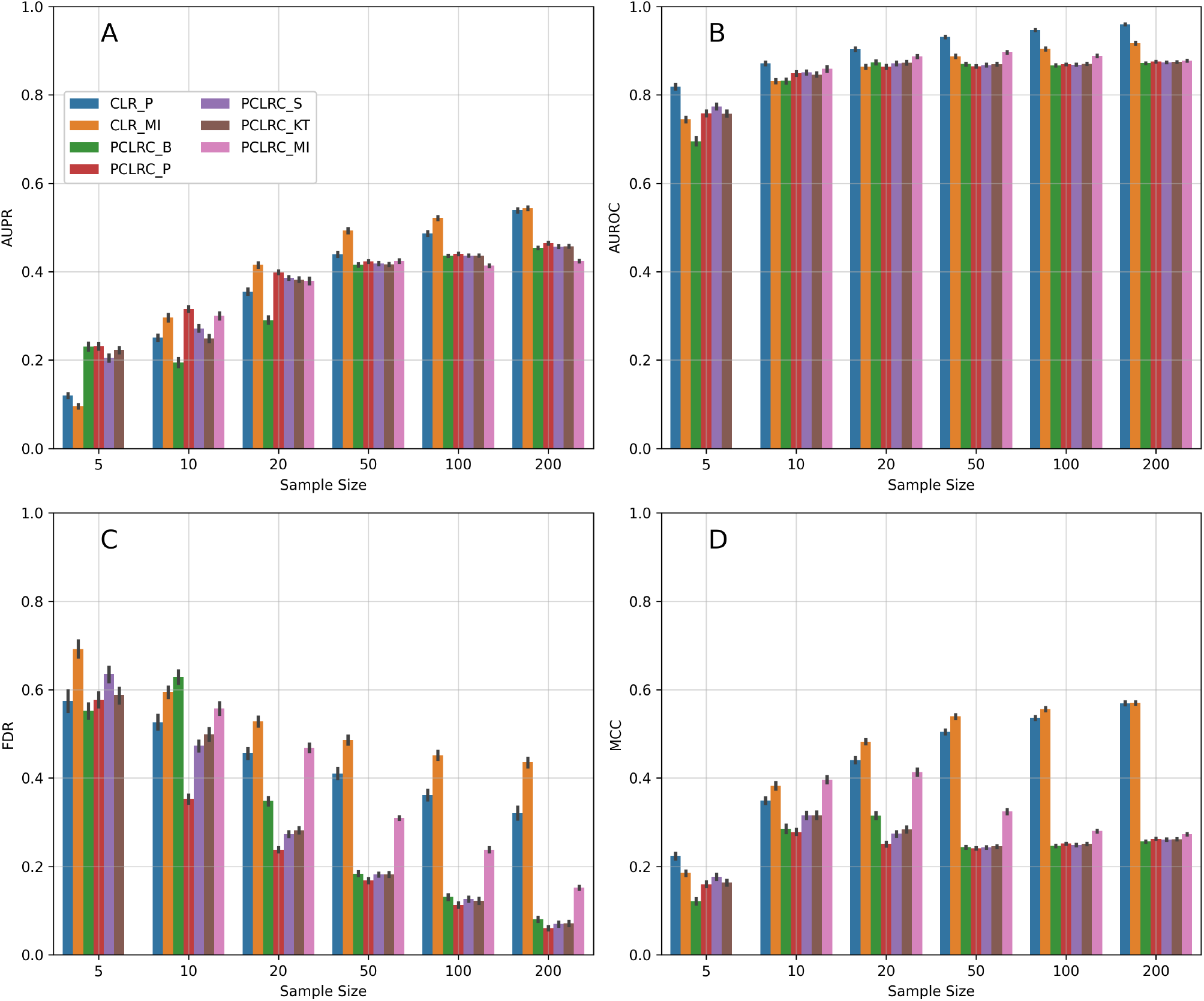
Network inference algorithms (NIAs) evaluated on pairwise performance measures. Thresholding values for FDR and MCC were calculated optimizing for F1-score for each individual network to limit the effect of threshold selection. This selection of NIAs are *CLR*-based methods for visual clarity. See Figure 6 for non-*CLR*-based methods.

**Supplementary Figure 8.**
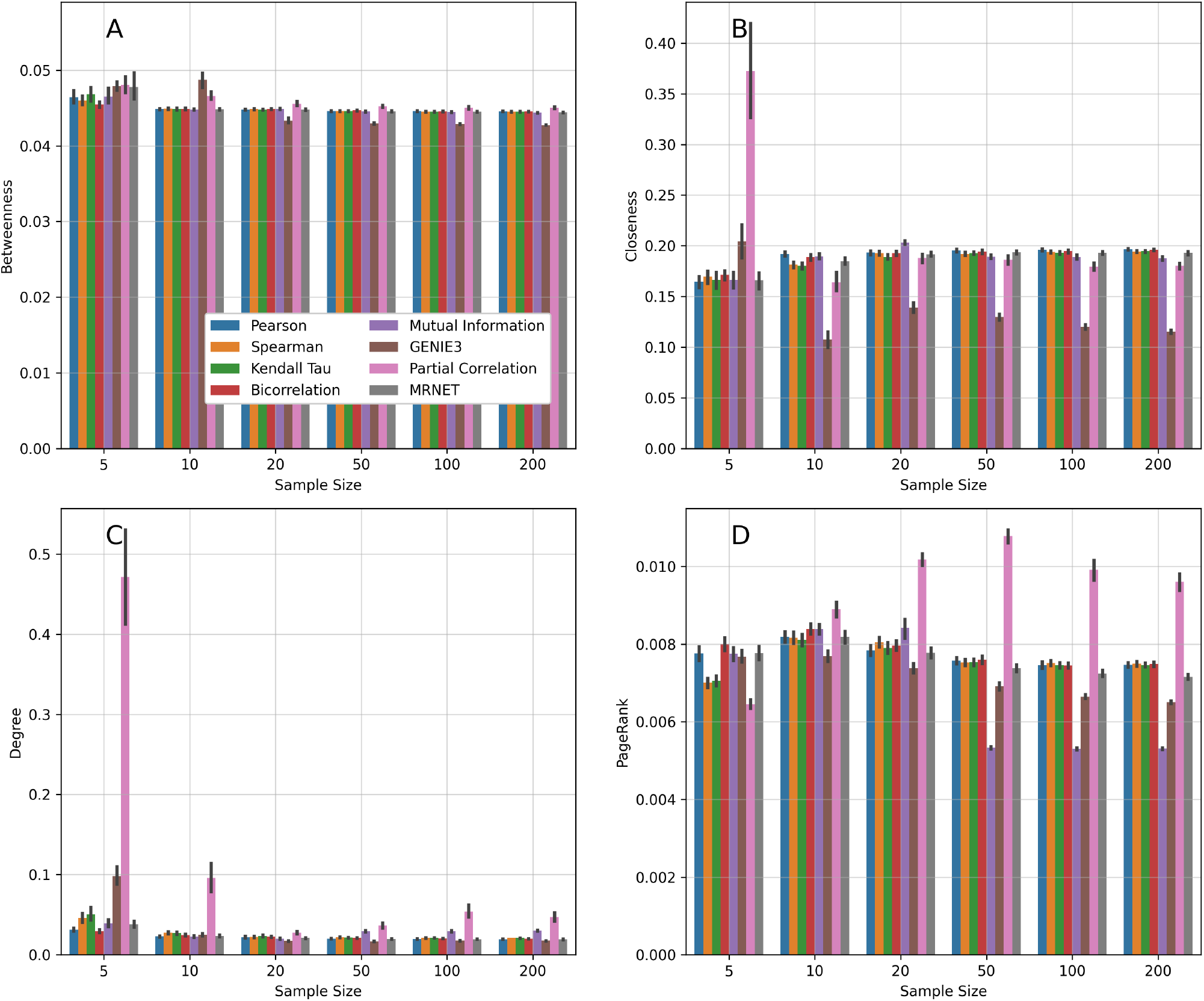
Network inference algorithms (NIAs) evaluated on network-level performance measures. Thresholding values were calculated optimizing for F1-score for each individual network to limit the effect of threshold selection. This selection of NIAs exclude *CLR*-based methods for visual clarity. See Figure 9 for *CLR*-based methods.

**Supplementary Figure 9.**
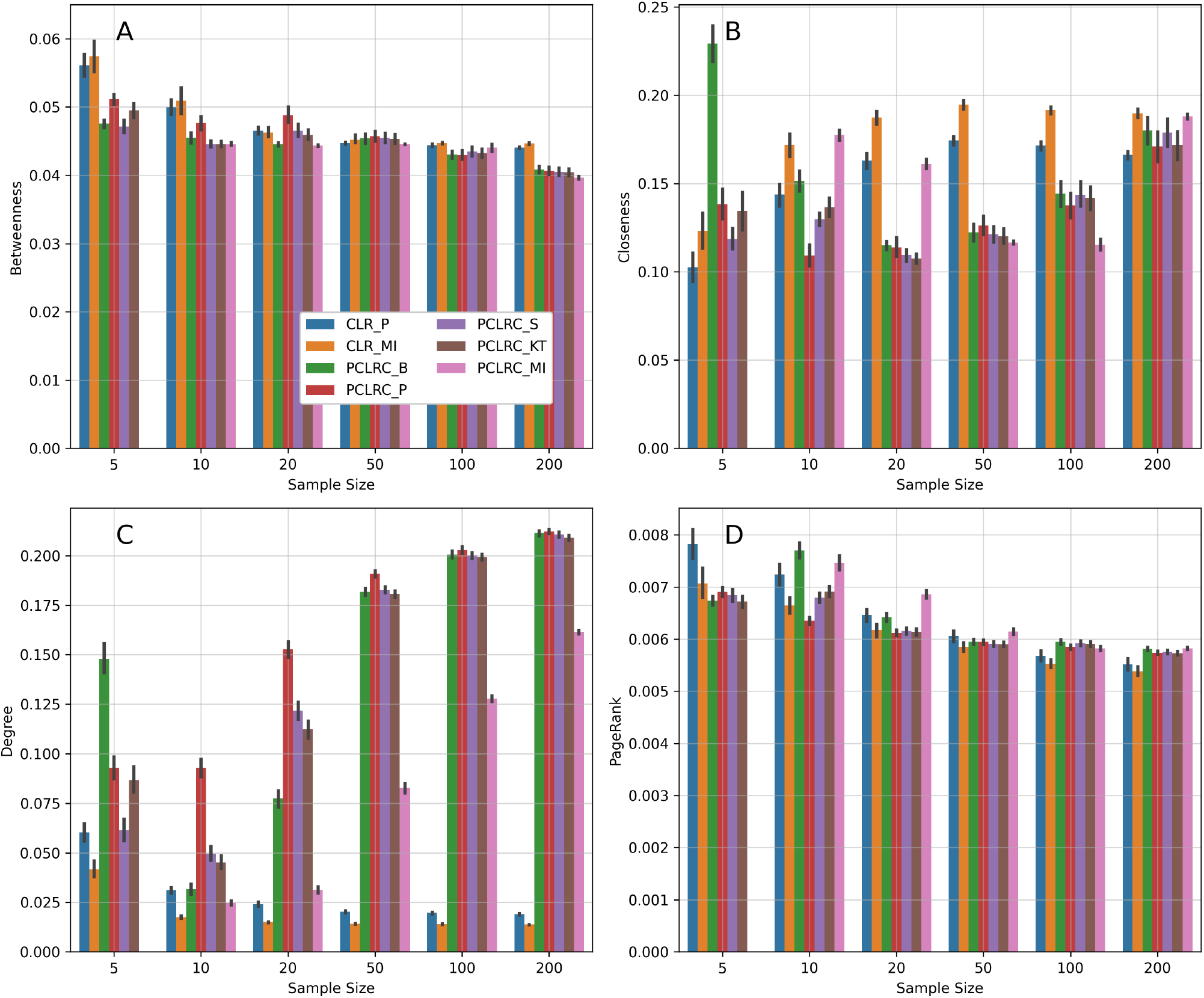
Network inference algorithms (NIAs) evaluated on network-level performance measures. Thresholding values were calculated optimizing for F1-score for each individual network to limit the effect of threshold selection. This selection of NIAs are *CLR*-based methods for visual clarity. See Figure 8 for non-*CLR*-based methods.

## References

S. Boughorbel, F. Jarray, and M. El-Anbari. Optimal classifier for imbalanced data using matthews correlation coefficient metric. PLOS ONE, 12, 2017.

C. Buccitelli and M. Selbach. mrnas, proteins and the emerging principles of gene expression control. Nature Reviews Genetics, 21:630 – 644, 2020.

A. J. Butte and I. S. Kohane. Mutual information relevance networks: functional genomic clustering using pairwise entropy measurements. Pacific Symposium on Biocomputing. Pacific Symposium on Biocomputing, pages 418–29, 1999.

D. Camacho, A. de la Fuente, and P. Mendes. The origin of correlations in metabolomics data. Metabolomics, 1:53–63, 2005.

D. Chicco and G. Jurman. The matthews correlation coefficient (mcc) should replace the roc auc as the standard metric for assessing binary classification. BioData Mining, 16, 2023.

S. de Siqueira Santos, D. Y. Takahashi, A. Nakata, and A. Fujita. A comparative study of statistical methods used to identify dependencies between gene expression signals. Brief Bioinform., 15:906–918, 2014.

C. Ding and H. Peng. Minimum redundancy feature selection from microarray gene expression data. Journal of Bioinformatics and Computational Biology, 3(2):185–205, 2005.

J. Faith, B. Hayete, J. Thaden, I. Mogno, J. Wierzbowski, G. Cottarel, S. Kasif, J. Collins, and T. Gardner. Large-scale mapping and validation of escherichia coli transcriptional regulation from a compendium of expression profiles. PLoS Biology, 5:e8, 2007.

L. Freeman. A set of measures of centrality based on betweenness. Sociometry, 40(1):35–41, 1977.

H. Greenyer. Evaluation of network inference algorithms and their effects on network analysis for the study of small metabolomic data sets. Master’s thesis, University of Victoria, 2022.

C. R. Harris, K. J. Millman, S. van der Walt, R. Gommers, P. Virtanen, D. Cournapeau, E. Wieser, J. Taylor, S. Berg, N. J. Smith, R. Kern, M. Picus, S. Hoyer, M. H. van Kerkwijk, M. Brett, A. Haldane, J.F. delR’io, M. Wiebe, P. Peterson, P. G’erard-Marchant, K. Sheppard, T. Reddy, W. Weckesser, H. Abbasi, C. Gohlke, and T. E. Oliphant. Array programming with numpy. Nature, 585:357 – 362, 2020.

S. Hoops, S. Sahle, R. Gauges, C. Lee, J. Pahle, N. Simus, M. Singhal, L. Xu, P. Mendes, and U. Kummer. Copasi—a complex pathway simulator. Bioinformatics, 22(24):3067– 3074, 2006.

V. A. Huynh-Thu, A. Irrthum, L. Wehenkel, and P. Geurts. Inferring regulatory networks from expression data using tree-based methods. PLOS ONE, 9(5), 2010.

S. Jahagirdar, M. Suarez-Diez, and E. Saccenti. Simulation and reconstruction of metabolite-metabolite association networks using a metabolic dynamic model and correlation based algorithms. Journal of Proteome Research, 18(3):1099–1113, 2019.

M. Kendall. A new measure of rank correlation. Biometrika, 1938.

J. Krumsiek, K. Suhre, T. Illig, J. Adamski, and F. J. Theis. Gaussian graphical modeling reconstructs pathway reactions from high-throughput metabolomics data. BMC Systems Biology, 5:21, 2011.

P.-J. Lahtvee, B. J. Sánchez, A. Smialowska, S. Kasvandik, I. E. Elsemman, F. Gatto, and J. B. Nielsen. Absolute quantification of protein and mrna abundances demonstrate variability in gene-specific translation efficiency in yeast. Cell systems, 4 5:495–504.e5, 2017.

S. K. Lam, A. Pitrou, and S. Seibert. Numba: a llvm-based python jit compiler. In LLVM ‘15, 2015.

J. Li, W. Weckwerth, and S. Waldherr. Network structure and fluctuation data improve inference of metabolic interaction strengths with the inverse jacobian. NPJ Systems Biology and Applications, 10, 2024.

C. Lingjærde, T. G. Lien, Ø. Borgan, H. Bergholtz, and I. K. Glad. Tailored graphical lasso for data integration in gene network reconstruction. BMC Bioinformatics, 22, 2020.

C. Liu, Y. Ma, J. Zhao, R. Nussinov, Y.-C. Zhang, F. Cheng, and Z.-K. Zhang. Computational network biology: Data, models, and applications. Physics Reports, 846:1–66, 2020.

N. Mahendran, P. M. D. R. Vincent, K. Srinivasan, and C.-Y. Chang. Improving the classification of alzheimer’s disease using hybrid gene selection pipeline and deep learning. Frontiers in Genetics, 12, 2021.

D. Marbach, J. C. Costello, R. Küffner, N. M. Vega, R. J. Prill, D. M. Camacho, K. R. Allison, M. Kellis, J. J. Collins, and G. Stolovitzky. Wisdom of crowds for robust gene network inference. Nature Methods, 9(8):796–804, 2012.

C. Marzban. The roc curve and the area under it as performance measures. Weather and Forecasting, 19(6): 1106–1114, 2004.

B. Matthews. Comparison of the predicted and observed secondary structure of t4 phage lysozyme. Biochimica et Biophysica Acta (BBA) - Protein Structure, 405(2): 442–451, 1975.

W. McKinney. Data structures for statistical computing in python. In SciPy, 2010.

P. Meyer, K. Kontos, F. Lafitte, and G. Bontempi. Information-theoretic inference of large transcriptional regulatory networks. EURASIP journal on bioinformatics & systems biology, 2007.

L. Michaelis, M. L. Menten, K. A. Johnson, and R. S. Goody. The original michaelis constant: translation of the 1913 michaelis-menten paper. Biochemistry, 50(39):8264–8269, 2011.

G. Nyamundanda, I. C. Gormley, Y. Fan, W. M. Gallagher, and L. Brennan. Metsizer: selecting the optimal sample size for metabolomic studies using an analysis based approach. BMC Bioinformatics, 14:338 – 338, 2013.

A. R. Palla, M. Ravichandran, Y. X. Wang, L. Alexandrova, A. Yang, P. E. Kraft, C. Holbrook, C. M. Schürch, A. T. V. Ho, and H. M. Blau. Inhibition of prostaglandin-degrading enzyme 15-pgdh rejuvenates aged muscle mass and strength. Science, 371, 2020.

K. Pearson. Notes on the history of correlation. Biometrika, 13(1):25–45, 1920.

F. Pedregosa, G. Varoquaux, A. Gramfort, V. Michel, B. Thirion, O. Grisel, M. Blondel, G. Louppe, P. Prettenhofer, R. Weiss, R. J. Weiss, J. Vanderplas, A. Passos, D. Cournapeau, M. Brucher, M. Perrot, and E. Duchesnay. Scikit-learn: Machine learning in python. ArXiv, abs/1201.0490, 2011.

L. Peel, T. P. Peixoto, and M. D. Domenico. Statistical inference links data and theory in network science. Nature Communications, 13, 2022.

E. Saccenti, M. Suarez-Diez, C. Luchinat, C. Santucci, and L. Tenori. Probabilistic networks of blood metabolites in healthy subjects as indicators of latent cardiovascular risk. Journal of Proteome Research, 14(2):1101–1111, 2015.

E. Saccenti, M. M. W. B. Hendriks, and A. K. Smilde. Corruption of the pearson correlation coefficient by measurement error and its estimation, bias, and correction under different error models. Scientific Reports, 10, 2019.

M. M. Saint-Antoine and A. Singh. Network inference in systems biology: recent developments, challenges, and applications. Current Opinion in Biotechnology, 63:89–98, 2020.

T. Saito and M. Rehmsmeier. The precision-recall plot is more informative than the roc plot when evaluating binary classifiers on imbalanced datasets. PLOS ONE, 10(3), 2015.

L. Song, P. Langfelder, and S. Horvath. Comparison of coexpression measures: mutual information, correlation, and model based indices. BMC Bioinformatics, 13(1):328, 2012.

C. Spearman. ‘general intelligence,’ objectively determined and measured. The American Journal of Psychology, 15(2):201– 293, 1904.

M. Suarez-Diez and E. Saccenti. Effects of sample size and dimensionality on the performance of four algorithms for inference of association networks in metabonomics. Journal of Proteome Research, 14:5119–5130, 2015.

J. Szymanski, S. Jozefczuk, Z. Nikoloski, J. Selbig, V. Nikiforova, G. Catchpole, and L. Willmitzer. Stability of metabolic correlations under changing environmental conditions in escherichia coli – a systems approach. PLOS ONE, 4(10):e7441, 2009.

R. Vallat. Pingouin: statistics in python. J. Open Source Softw., 3:1026, 2018.

P. Virtanen, R. Gommers, T. E. Oliphant, M. Haberland, T. Reddy, D. Cournapeau, E. Burovski, P. Peterson, W. Weckesser, J. Bright, S. J. van der Walt, M. Brett, J. Wilson, K. J. Millman, N. Mayorov, A. R. J. Nelson, E. Jones, R. Kern, E. Larson, C. J. Carey, İlhan Polat, Y. Feng, E. W. Moore, J. Vanderplas, D. Laxalde, J. Perktold, R. Cimrman, I. Henriksen, E. A. Quintero, C. R. Harris, A. M. Archibald, A. H. Ribeiro, F. Pedregosa, P. van Mulbregt, A. A. P. A. A. A. A. A. V. B. R. H. K. Sco, A. Vijaykumar, A. P. Bardelli, A. Rothberg, A. Hilboll, A. Kloeckner, A. M. Scopatz, A. Lee, A. S. Rokem, C. N. Woods, C. Fulton, C. Masson, C. Häggström, C. Fitzgerald, D. A. Nicholson, D. R. Hagen, D. V. Pasechnik, E. Olivetti, E. Martin, E. Wieser, F. Silva, F. Lenders, F. Wilhelm, G. Young, G. A. Price, G.-L. Ingold, G. E. Allen, G. R. Lee, H. Audren, I. Probst, J. P. Dietrich, J. Silterra, J. T. Webber, J. Slavić, J. Nothman, J. Buchner, J. Kulick, J. L. Schónberger, J. V. D. M. Cardoso, J. Reimer, J. E. Harrington, J. Rodríguez, J. Nunez-Iglesias, J. Kuczynski, K. L. Tritz, M. Thoma, M. Newville, M. Kümmerer, M. Bolingbroke, M. Tartre, M. Pak, N. J. Smith, N. Nowaczyk, N. Shebanov, O. Pavlyk, P. A. Brodtkorb, P. Lee, R. T. McGibbon, R. Feldbauer, S. Lewis, Tygier, S. Sievert, S. Vigna, S. Peterson, S. More, Pudlik, T. Oshima, T. J. Pingel, T. P. Robitaille, T. Spura, T. R. Jones, T. Cera, T. Leslie, T. Zito, T. Krauss, U. Upadhyay, Y. O. Halchenko, and Y. Vázquez-Baeza. Scipy 1.0: fundamental algorithms for scientific computing in python. Nature Methods, 17:261 – 272, 2019.

J. M. Wilkins and E. Trushina. Application of metabolomics in alzheimer’s disease. Frontiers in Neurology, 8:719, 2018.

D. Wishart. Current progress in computational metabolomics. Briefings Bioinform., 2007.

C.-H. Zheng, L. Yuan, W. Sha, and Z.-L. Sun. Gene differential coexpression analysis based on biweight correlation and maximum clique. BMC Bioinformatics, 15, 2015.

